# Spatiotemporal and single-cell atlases to dissect cell lineage differentiation and regional specific cell types in mouse ovary morphogenesis

**DOI:** 10.1101/2023.07.21.549985

**Authors:** Zheng-Hui Zhao, Tie-Gang Meng, Fei Gao, Heide Schatten, Qing-Yuan Sun

## Abstract

Characterization of cell heterogeneity and molecular diversity using single-cell RNA sequencing has greatly enhanced our understanding of the ovary’s dynamic differentiation processes. However, regional specification of ovarian cells for certain physiological functions remains largely unexplored in the physical space. Here, we combine spatial transcriptomics with single-cell RNA sequencing technologies to build a spatiotemporal and single-cell atlas of ovaries from fetal to adult stages. We construct the pseudotime trajectories of female germ cells and bipotential pregranulosa cells and define key regulatory transcription factors responsible for their differentiation processes. Specifically, we dissect the relationships between two waves of meiosis initiation, oogenesis processes and folliculogenesis. Moreover, we characterize the region-specific subtypes of granulosa cells and luteal cells and construct pseudo-space-time trajectories from granulosa cells to luteal cells. Notably, we identify small luteal cells, a novel cell type, which highly express *Onecut2* and exclusively locate at the corpus luteum. Altogether, this study comprehensively delineates ovary development and regional specific ovarian cell types.

## Introduction

In mammals, somatic cells of the ovary are derived from the genital ridge that forms as a thickening of the coelomic epithelium on the ventromedial surface of the mesonephros. After localizing to the genital ridge, primordial germ cells proliferate actively with incomplete cytokinesis to form germ cell nests [1]. Subsequently, activation of Rspo1/Wnt4/β-catenin signaling pathway promotes *Foxl2* expression and granulosa cell differentiation [2]. Female germ cells proceed mitosis to meiosis transition directly following granulosa cell differentiation and then arrest at the diplotene stage of prophase I [3]. Around the period of follicle assembly, the germ cell nests in the medullary region begin to break down to form the first wave of follicles that grow directly without activation, and then the breakdown of germ cell nests in the cortical region occurs to form the primordial follicles that grow after recruitment and activation [4]. With the completion of follicle assembly, a well-organized ovary with complex cell types forms. As the most dynamic organ, the ovary undergoes dramatic restructuring for the process of the menstrual cycle. After the first wave of folliculogenesis, the follicles are recruited to the cortex, start to grow in the medulla and finally ovulate at the ovarian surface. After ovulation, the basement membrane of follicles undergo breakdown, the microvascular cells invade granulosa layers and the luteinizing granulosa cells and theca cells encroach on the follicular cavity, which subsequently forms the corpus luteum [5]. The corpus luteum is a temporary endocrine structure in the ovary, which mainly includes small luteal cells, large luteal cells, endothelial cells and immune-related cells. Of these cell types, the small luteal cells and large luteal cells, which derive from theca cells and granulosa cells, respectively, are capable of secreting progesterone that is critical for maintaining pregnancy [6]. However, the majority of ovarian cell types as well as their spatial distribution are incompletely characterized and the molecular diversity and cell fate determination of ovarian cells remain elusive, suggesting the necessity of using single-cell approaches to decode the dynamic changes of cell types at the molecular and spatial levels.

Recent advances in single-cell RNA sequencing (scRNA-seq) have enabled the comprehensive illustration of molecular heterogeneity at the single-cell resolution. A comprehensive and systematic view of multi-lineage specification and gene-regulatory modules may help us to understand ovary differentiation and maturation. Work from several groups has explored the ovary development and pathology in different species [4, 7–15]. In previous studies, researchers only detected fetal and early postnatal development of the mouse ovary [4, 7–10], while the later development of the ovary has not yet been explored. Although scRNA-seq approaches can profile the transcriptomes at the single-cell level, the spatial information of cells cannot be preserved. As emerging technologies, spatial transcriptomics provide new opportunities to profile spatial regions in tissue sections and define the organization of cellular niches [16–18]. At present, several researches have applied these excellent technologies to investigate human gonadal development [19] and mouse ovary aging [20] at certain developmental stages. However, no work could profile the complete spatiotemporal development of the mouse ovary at the single cell level.

Here, by using scRNA-seq and spatial transcriptional profiling, we investigate the cell types and spatial distribution of the mouse developing ovary. Several computational methods are applied in this study to resolve the spatial relationships of various cell types involved in follicle development and we explore previously uncharacterized cell types based on their transcriptional features and locations in the original tissue sections. In the present study, we identified 11 cell types and constructed developmental trajectories from the early undifferentiated gonad to the mature adult ovary. We dissected region-specific granulosa cells in different ovary developmental stages. For example, cumulus granulosa cells (cGC) and mural granulosa cells (mGC) in different regions of the follicle varied dramatically in expression patterns and ratios. Additionally, the distribution of luteal cells exhibited regional specificities, and the large luteal cells (LLC) locate both in the corpus luteum and other regions of the ovary, while the small luteal cells (SLC) that highly express *Onecut2* exclusively locate in the corpus luteum, which lays the foundation of the functional significance in the corpus luteum. Among them, what is exciting is that we revealed the molecular diversity and spatial distribution of luteal cells in the adult ovary for the first time, which may regulate the secretion of progesterone in the corpus luteum. We anticipate that our results will enhance our current understanding of different cell types in the corpus luteum and provide directions for potential treatment of progesterone-related subfertility.

## Results

### Single-cell and spatiotemporal atlas of the developing mouse ovary

To understand the dynamic interactions between differentiation and morphogenesis of the mouse ovary, we performed single-cell and spatial transcriptomics. We profiled 50,655 single cells from twelve developmental stages ranging from the early undifferentiated gonad (embryonic day 11.5 gonad, E11.5 gonad) to the mature adult ovary (postnatal day 90 ovary, PD90 ovary) using the 10X Genomics scRNA-seq technique. The previously published data represented approximately 75% of the cells in the single-cell dataset [4, 8, 9], and the single cell data of PD0, PD21 and PD90 has been sequenced in this study (Fig. 1A). After quality control, batch effect correction with Harmony [21], and graph-based clustering with Seurat [22], we obtained 11 transcriptionally distinct cell populations (Fig. 1B). Based on the classic marker genes, we annotated them as female germ cells (FGCs; *Ddx4*, *Dazl*), bipotential pregranulosa cells (BPG; *Foxl2*, *Fst*), epithelial pregranulosa cells and epithelial cells (EPG&Epi; *Lgr5*, *Lhx9*, *Amhr2*), early theca (*Dlk1*, *Coloa1*), theca (*Hsd3b1*, *Cyp11a1*), proliferative mesenchyma (pMesenchyma; *Pclaf*, *Lhx9*), mesenchyma (*Col1a1*, *Nr2f2*), endothelial cells (Endo; *Pecam1*, *Cldn5*), pericyte (*Rgs5*), immune cells (*Tyrobp*), erythrocyte (*Alas2*) (Fig. 1C and fig. S1A). Additionally, gene ontology (GO) analysis of the top 200 differentially expressed genes (DEGs) across cell types revealed features corresponding to known biological functions and characteristics of each cell cluster (fig. S1B). For instance, the GO terms “retinoic acid binding” and “adrenergic receptor binding” were specific to the early theca cell cluster, whereas “steroid dehydrogenase activity” and “cholesterol transfer activity” were specific to the theca cell cluster (fig. S1B). We further quantified the relative proportions of these cell types at different ovary developmental stages and examined the distribution of these cell clusters in each donor sample (Fig. 1D and E). These cell types varied significantly in global relatedness according to the expression patterns of known marker genes and GO terms. We then turned our attention to dynamic changes and the transition of several key cell types associated with meiosis initiation, follicle assembly as well as development and corpus luteum formation, including female germ cells, granulosa cells and theca cells during ovary morphogenesis. We found that these clusters changed in a development-dependent manner.

**Figure 1.**
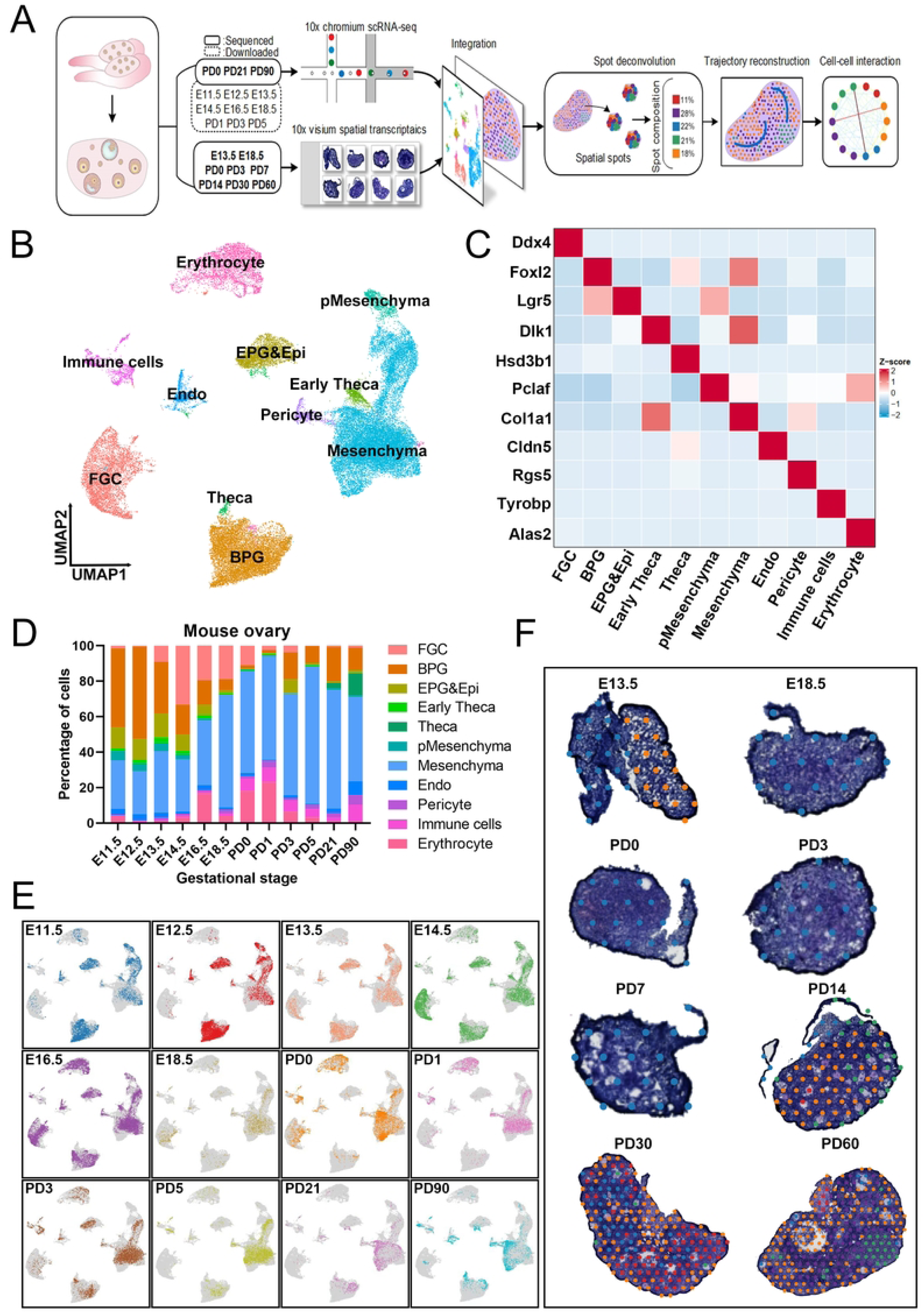
Spatially resolved single-cell transcriptomic landscape of the developing mouse ovary. (A) Schematic diagram of the experimental workflow and analysis for single-cell RNA-seq and spatial transcriptomic data. (B) UMAP projection of 50,655 cells (divided into 11 clusters) from 12 pooled samples of ovaries, colored by cell type. BPG, bipotential pregranulosa cells; Endo, endothelial cells; EPG&Epi, epithelial pregranulosa cells and epithelial cells; FGC, female germ cells; pMesenchyma, proliferative mesenchyma. (C) Heatmap of cell type-specific marker genes. (D) The percentage of ovarian cells classified by developmental stages. (E) UMAP projection of single-cell transcriptomes colored by developmental stage from E11.5 to PD 90. E11.5, embryonic day 11.5; PD90, postnatal day 90. (F) Spatial spots clustered by gene expression and labeled by tissue anatomical compartment for eight developmental stages.

Spatial organization is essential for understanding the physiological functions of various ovarian cell types and the signaling interactions between them. To precisely decode spatiotemporal ovary developmental events, we performed spatial transcriptomic analysis on mice at eight developmental stages ranging from the early fetal gonad (E13.5 gonad) to the mature adult ovary (PD60 ovary). After mapping and filtering, we obtained 1,120 individual spots and 5483 median genes for each spot. First, we performed clustering on stSME normalized spatial transcriptomics data using stLearn [23] and identified different clusters on the ovary sections after PD14, which indicates that the early ovaries exhibit relative uniform distribution of unique transcripts (Fig. 1F and fig. S1C). Subsequently, we also examined the spatial expression patterns of canonical marker genes for female germ cells, granulosa cells and theca cells, which reveals a well-organized distribution pattern composed of distinct cell clusters (fig. S1D). In summary, using spatial transcriptomic profiling combined with single-cell RNA sequencing, we described a comprehensive development atlas of the mouse ovary.

### Characteristics of two waves of oogenesis in the developing mouse ovary

In the previous study, we have depicted two waves of oogenesis in fetal monkey FGCs [11]. However, the pseudotime trajectory of two-wave oogenesis in mouse FGCs has not been constructed owing to limited transcriptome data. Here, we integrated the scRNA-seq data of FGCs from E11.5 to PD5 developmental stages and divided them into seven clusters (Fig. 2A). To dissect the heterogeneity of FGCs in the early postnatal days, we examined the distributions of FGCs in PD0, PD1, PD3 and PD5 stages; the PD0 stage mainly contains growing and dormant oocytes, while the PD1, PD3 and PD5 stages mainly include growing oocytes (fig. S2A). To further identify the subtypes of FGCs, we characterized the stage-specific marker genes for FGC subtypes; the mitotic FGCs (oogonia and pre-meiotic oogonia) highly expressed pluripotent genes, such as *Nanog*, *Dppa5a* and *Pou5f1*, while meiotic FGCs highly expressed meiotic marker genes, such as *Spo11*, *Prdm9* and *Sycp3* (Fig. 2b; Supplementary Fig. S2b). Notably, the dormant oocytes highly expressed *S100a9* and *S100a8*, while the growing oocytes highly expressed *Figla*, *Sohlh1* and *Nobox*, which suggests that these genes may participate in the follicle dormancy or activation (Fig. 2B).

**Figure 2.**
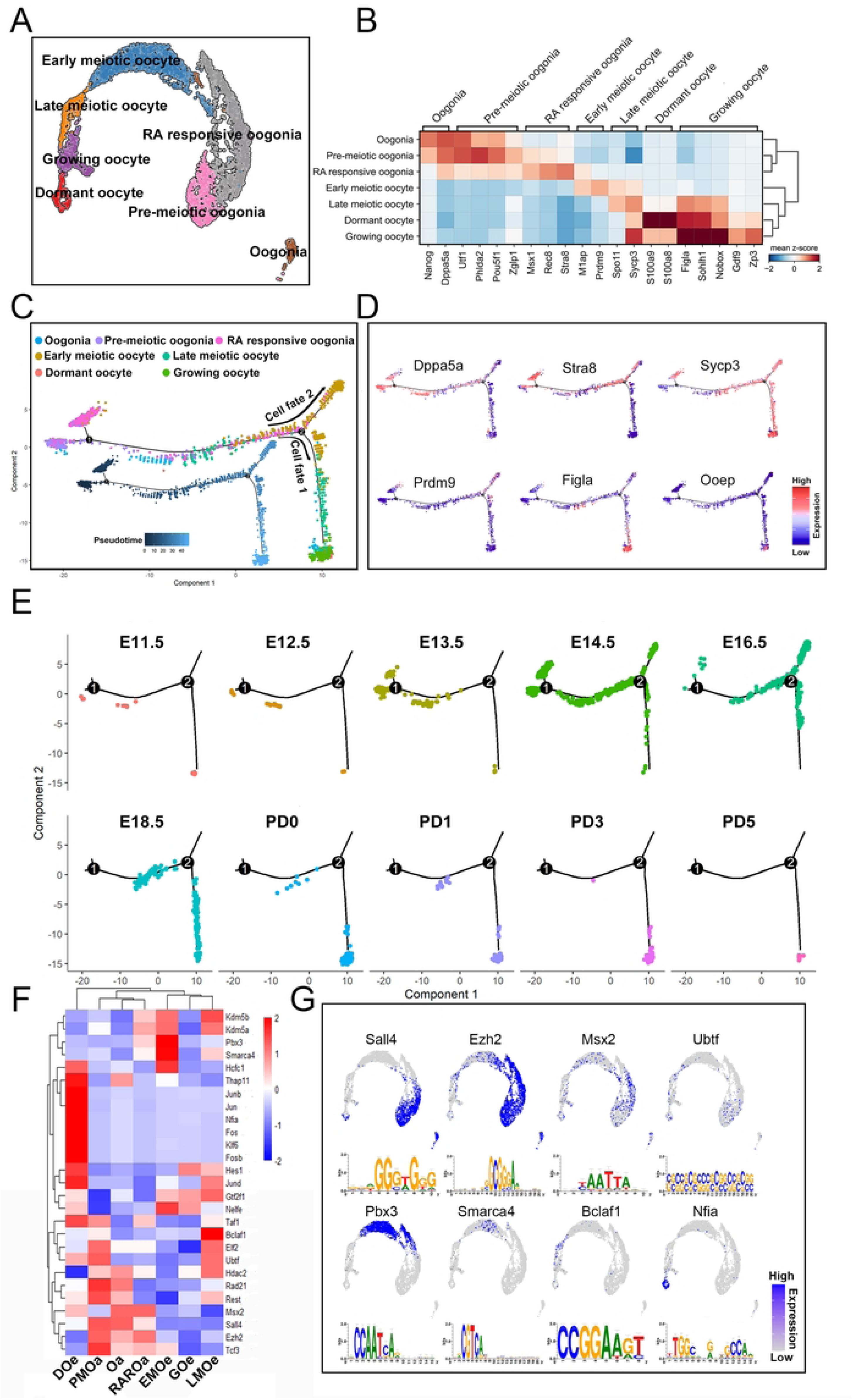
Molecular diversity and developmental trajectory of female germ cells. (A) Visualization of FGC subclusters using UMAP. RA, retinoic acid. (B) Heatmap showing the marker genes of each female germ cell subcluster. (C) Pseudotime trajectory of female germ cells analyzed by Monocle. (D) Expression of representative genes along pseudotime trajectory. (E) Pseudotime trajectory of female germ cells split by developmental stages. (F) Heatmap of regulon (the regulatory network of transcription factors and their target genes) activity analyzed by SCENIC. Doe, dormant oocyte; EMOe, early meiotic oocyte; GOe, growing oocyte; LMOe, late meiotic oocyte; Oa, oogonia; PMOa, pre-meiotic oogonia; RAROa, RA responsive oogonia. (G) UMAP showing representative regulons and their first targeted motifs.

We next constructed the pseudotime trajectory of FGCs using monocle, which represented two waves of oogenesis (Fig. 2C). The first wave of oogenesis occurs on the branch 1 around meiosis initiation and the second wave of oogenesis occurs on the branch 2 around follicle assembly. Meanwhile, the representative genes, such as *Prdm9*, *Figla* and *Ooep*, are displayed along the pseudotime trajectory, which indicates that the FGCs in cell fate 1 may survive to form follicles, while the majority of early meiotic oocytes in cell fate 2 may undergo germ cell attrition (Fig. 2C and D). To further explore the cell fate determination on the branch 2, two gene sets with distinct patterns were identified, which were likely involved in the determination of female germ cell fates. The set 1 includes genes, such as *Sohlh1*, *Uchl1* and *Xdh,* that are highly expressed in follicular oocytes, while the set 2 includes genes, such as *Inca1*, *Stag3* and *Pbx3,* that are highly expressed in dying oocytes (fig. S2C to E). To dissect the occurrence time of two waves of oogenesis, we presented the pseudotime trajectory with facet plots according to the developmental stages, which showed that the first wave of oogenesis occurs at E13.5 and the second wave of oogenesis mainly occurs at E16.5 (Fig. 2E). Interestingly, in the early developmental stages from E11.5 to E14.5, a few germ cells undergo the second wave of oogenesis (Fig. 2E), which may serve as pioneer cells to monitor the process of second oogenesis.

To explore the regulatory mechanisms underlying the FGC status transition, we performed single-cell regulatory network inference and clustering (SCENIC) and DorothEA analysis and discovered that dozens of transcription factors displayed a dynamic pattern. Hierarchical clustering analysis on the regulons (transcription factors and their targets) also revealed the regulatory relationship among the different FGC subclusters (Fig. 2F). For example, both of SCENIC and DorothEA revealed that dormant oocytes (DOe) displayed a unique regulons activity, and *Thap11*, *Jun* and *Jund* were active mainly in the DOe cluster (Fig. 2F and fig. S2F). Additionally, *Sall4*, *Pbx3*, *Bclaf1* and *Nfia* seemed to be mostly activated in the oogonia (Oa), early meiotic oocytes (EMOe), late meiotic oocytes (LMOe) and DOe, respectively (Fig. 2F). Moreover, these representative regulons were also mapped to uniform manifold approximation and projection (UMAP) plots and their motifs were listed (Fig. 2G). Furthermore, a portion of transcription factors that are solely enriched in subclusters appeared to serve a specific role, such as *Pbx3* may have a limited role in oocyte attrition and *Nfia* may function to maintain follicle dormancy. Taken together, our data depicted two waves of oogenesis that may be associated with follicle growth in ovary medulla and follicle dormancy in the ovary cortex.

### Molecular developmental trajectory of bipotential pregranulosa cells

After sex determination, a portion of ovarian somatic cells will differentiate into BPG which expresses *Foxl2* or EPG which expresses *Lgr5* and then expresses *Foxl2* robustly. In this study, the EPG and Epi were grouped into one cluster (EPG&Epi in Fig. 1B) and the EPG expresses *Foxl2* robustly after expressing *Lgr5*. To avoid the impact of Epi, the BPG cluster was therefore extracted and then divided into nine subtypes of granulosa cells (Fig. 3A) and the cells generated from the different developmental stages were labeled according to their derivation (Fig. 3B). To understand the molecular features of the BPG subtypes, we examined the typical marker genes for different subtypes. Nearly all BPG subtypes express *Kctd14* and *Prdx1* except for mGC that highly expresses *Prss23* and *Grb14* (Fig. 3C and fig. S3A). Additionally, the ribosomal protein genes associated cortical pregranulosa cells (Rp-CPG) and EPG highly express cortical marker genes, such as *Gng13* and *Lgr5*, suggesting that these cells may locate at the ovarian surface; while the medullary pregranulosa cells (MPG) and BPG highly express medullary marker genes, such as *Hmgcs2* and *Foxl2*, suggesting that these cells may locate at the ovarian center (Fig. 3C and fig. S3A).

**Figure 3.**
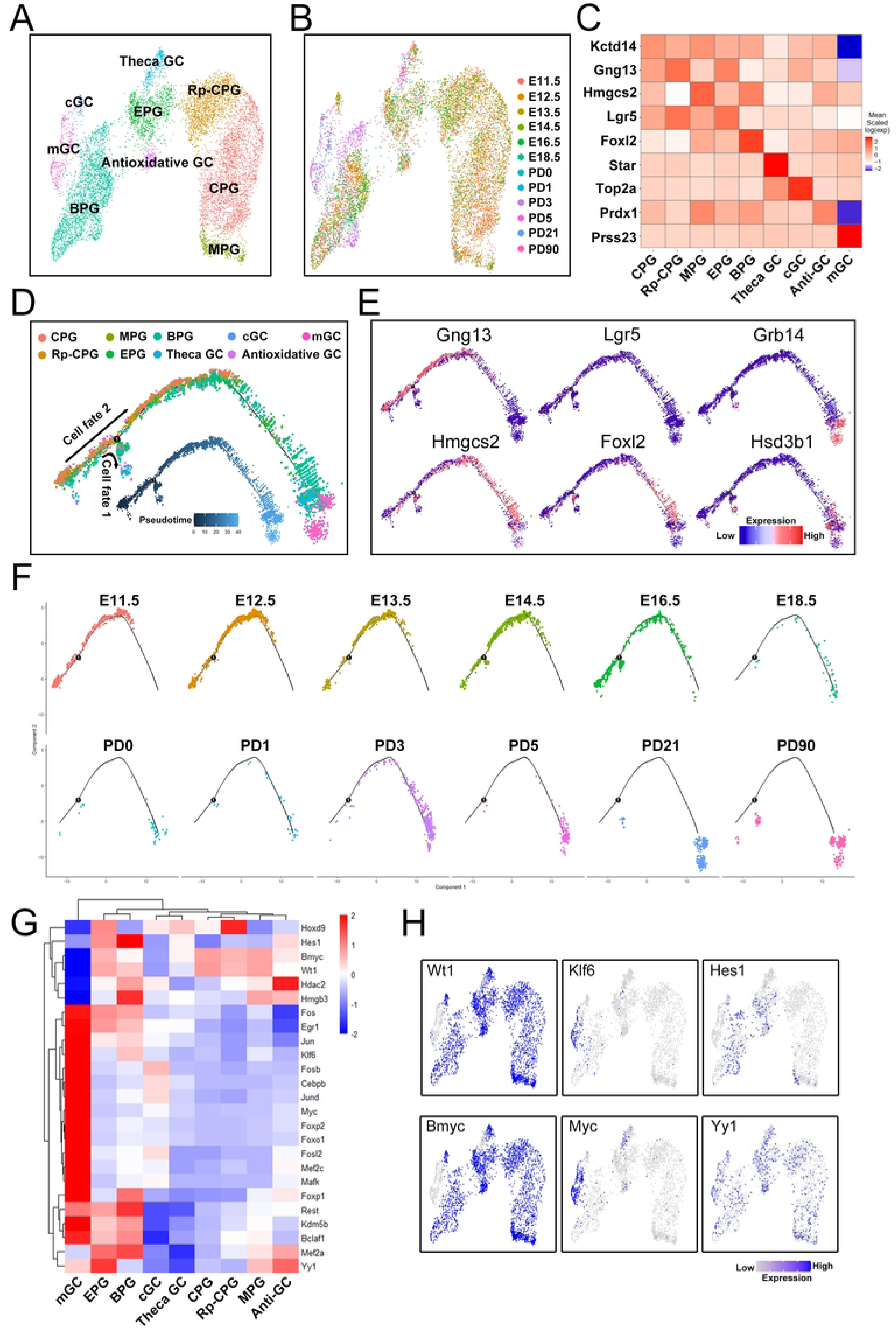
Molecular signatures and pseudotime trajectory of granulosa cells. (A) Visualization of major subtypes of BPG using UMAP. Antioxidative GC, antioxidative granulosa cells; BPG, bipotential pregranulosa cells; cGC, cumulus granulosa cells; CPG, cortical pregranulosa cells; EPG, epithelial pregranulosa cells; MPG, medullary pregranulosa cells; mGC, mural granulosa cells; Rp-CPG, ribosomal protein genes associated cortical pregranulosa cells; Theca GC, granulosa cells that differentiated into Theca cells. (B) UMAP projection of subtypes of BPG colored by developmental stages. (C) Heatmap of marker genes in each subtype of BPG. Anti-GC, antioxidative granulosa cells. (D) Pseudotime trajectory of bipotential pregranulosa cells analyzed by Monocle. (E) Expression of marker genes along pseudotime trajectory. (F) Pseudotime trajectory of bipotential pregranulosa cells split by developmental stages. (G) Heatmap representing the of activities of regulons analyzed by SCENIC. Anti-GC, antioxidative granulosa cells. (H) UMAP projection of representative regulons.

To explore the cell type transition and differentiation of BPG at the molecular level, we performed pseudotime analysis using monocle to construct the developmental trajectory (Fig. 3D). In the trajectory, cells progressed in pseudotime according to the differentiation state characterized by known marker genes, such as *Hmgcs2*, *Foxl2*, *Lgr5* and *Grb14* (Fig. 3E). Additionally, the deconvoluted pseudotime plot showed the branch 1 that indicates that the BPG undergo two cell-fate determinations (Fig. 3D). To further dissect the cell fate determination of BPG on the branch 1, two gene sets with distinct patterns were identified. The set 1 includes genes, such as *Hsd17b1*, *Serpine2* and *Cebpb,* that are highly expressed in granulosa cells that will differentiate into theca cells (Theca GC), while the set 2 includes genes, such as *Tsc22d1*, *Maged2*, *Foxl2* and *Col3a1*, that are highly expressed in mGC and Theca GC (fig. S3B and C). To dissect the time for BPG cell fate determination, we split the pseudotime trajectory with facet plots according to the developmental stages and found that the cell fate determination of Theca GC mainly occurs at E16.5, which is consistent with the second wave of oogenesis (Fig. 2E and Fig. 3F). The relationship between Theca GC and the second wave of oogenesis needs further exploration.

To understand the regulatory network underlying the BPG differentiation, we performed SCENIC and DorothEA analysis on BPG subtypes and discovered a series of transcription factors with a dynamic pattern. Hierarchical clustering analysis on the regulons revealed the regulatory relationship among the different BPG subclusters (Fig. 3G). For instance, both of SCENIC and DorothEA revealed that mGC displayed a unique regulons activity, and *Fosl2*, *Foxp1* and *Foxo1* were active mainly in the mGC cluster (Fig. 3G and fig. S3D). Moreover, several representative regulons were also mapped to UMAP plots and the *Wt1* and *Bmyc* were highly expressed in cells except for the mGC cluster, while the *Klf6* and *Myc* were highly expressed in the mGC cluster (Fig. 3H). Collectively, we revealed that the majority of *Foxl2* positive cells and a few *Lgr5* positive cells could differentiate into theca cells mainly at the E16.5 or postnatal stages.

### Two waves of meiosis initiation supporting two waves of folliculogenesis

The traditional meiosis initiation (second wave) occurs asynchronously from the ovary anterior to the posterior region [24]. Recently, it was shown that an earlier meiosis initiation (first wave) occurs with a radial manner from the ovary medial to the surface region [25]. To dissect the meiosis initiation processes, we examined the spatial distribution of pluripotent and meiotic associated genes enhanced by BayesSpace [26] (Fig. 4A and fig. S3E and F). Specifically, we revealed that *Zglp1*, a transcriptional regulator essential for meiotic entry [27], was highly expressed in two regions of the E13.5 gonad, and the *Cdx2*, a transcription factor required for repression of *Pou5f1* [28], was highly expressed around the *Zglp1*, which was further confirmed by immunostaining on the E13.5 gonad sections (Fig. 4A and B). Both, *Zglp1* and *Cdx2,* could promote the mitosis to meiosis transition through upregulation of meiotic genes and downregulation of pluripotent genes. Moreover, we could not detect the *Zglp1* expression on the ovary surface sections, suggesting that the first wave of meiotic entry may occur at the ovary medial region (fig. S3E and F). Additionally, the meiotic genes, such as *Msx1*, *Stra8*, *Sycp3* and *Rec8*, are expressed in a multicenter manner (Fig. 4A and fig. S3E and F), suggesting that the first wave of meiosis initiation occurs simultaneously in different ovary regions.

**Figure 4.**
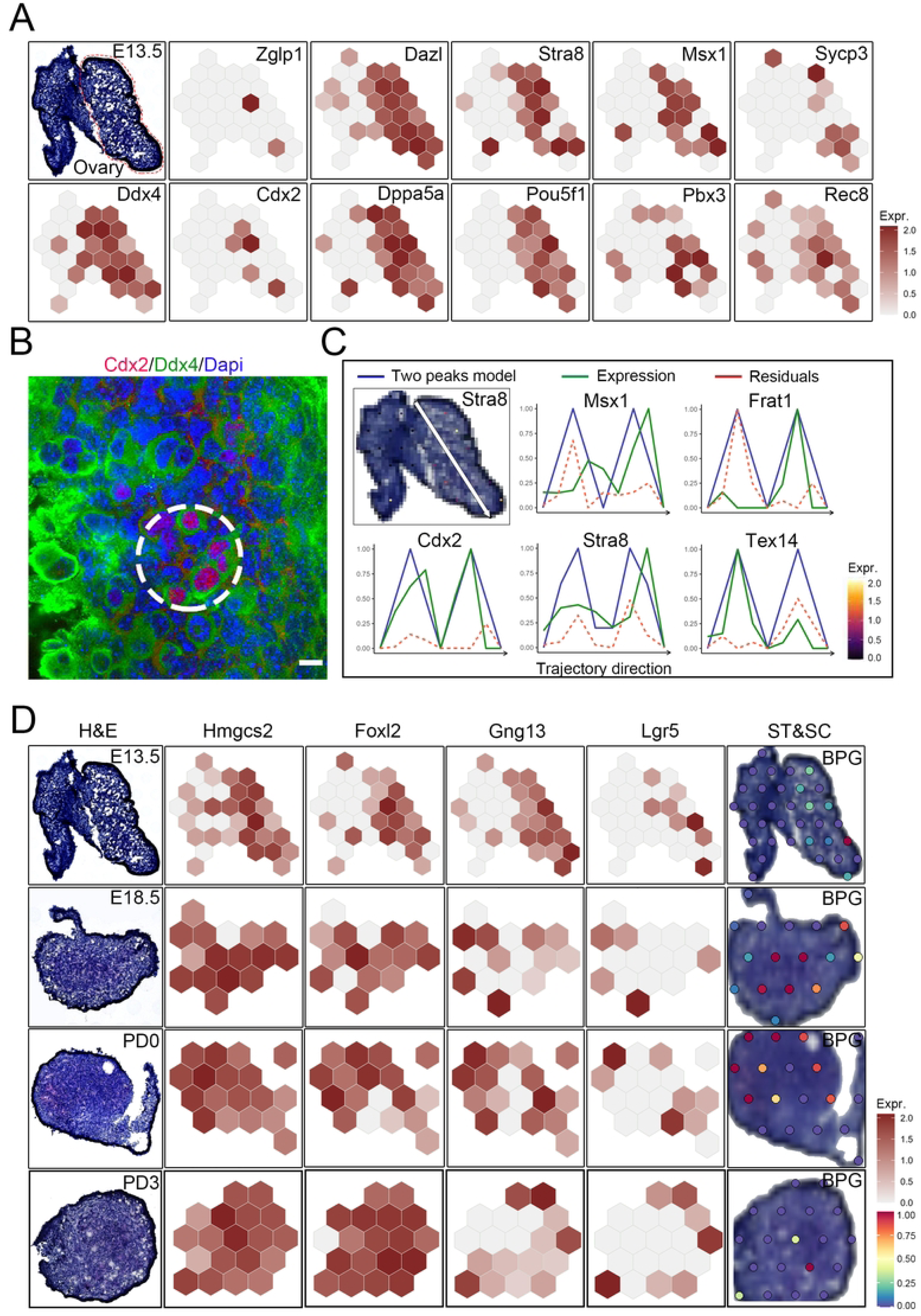
Spatial transcriptomics identify spatially restricted genes during the transition from mitosis to meiosis and follicle formation. (A) Spatially resolved gene expression for meiosis initiation associated genes enhanced by BayesSpace. (B) Immunostaining for Cdx2 and Ddx4 in mouse ovary. (C) Visualization of gene expression along the trajectory from anterior to posterior gonad analyzed by SPATA. (D) Spatially resolved gene expression for cortical and medullary follicle formation. H&E, hematoxylin and eosin; ST&SC, spatial transcriptomics and single cell transcriptomes.

To understand the second wave of meiosis initiation, we constructed a spatial trajectory (white arrow in Fig. 4C) from the ovary anterior to the posterior region and examined the expression patterns of pluripotent or meiotic genes using SPATA [29]. The expression patterns of several meiotic genes, such as *Msx1*, *Stra8*, *M1ap* and *Sycp3* exhibit two peaks along the spatial trajectory, and the former expression peak is low compared to the latter expression peak, suggesting that meiosis in the ovary anterior region has occurred and the relevant meiotic genes were downregulated (Fig. 4C and fig. S4A). Moreover, the spatial distribution of *Stra8* and *Tex14*, an intercellular bridge component required for meiotic entry within a cyst, is more spread in the anterior region than that in the posterior region (fig. S4B). Furthermore, we integrated the single-cell and spatial transcriptomic data of the E13.5 ovary and mapped the mitotic FGCs and meiotic FGC1 to the spatial section using Seurat, which revealed that the meiotic FGC1 mainly located at the anterior region, while the mitotic FGCs mainly located at the posterior region of the ovary (fig. S4B).

To further explore the relationship between two waves of meiosis and folliculogenesis, we identified the BPG and EPG clusters at E13.5, E18.5, PD0 and PD3 developmental stages using known marker genes, such as *Hmgcs2*, *Foxl2*, *Gng13* and *Lgr5* (fig. S4C). Moreover, the spatial expression patterns of these marker genes were also examined; the *Hmgcs2* and *Foxl2* are highly expressed in the ovary medulla, meanwhile, *Gng13* and *Lgr5* are highly expressed in the ovary cortex (Fig. 4D and fig. S4D). Furthermore, we mapped the BPG cluster in each stage to the relevant spatial sections and found that the BPG cluster mainly located at the ovary medulla at the E18.5 stage, and subsequently located at the ovary cortex at the PD0 stage, suggesting that cortical granulosa cells begin to express *Foxl2* and further form primordial follicles (Fig. 4D). Altogether, these data revealed that the medullary germ cells undergone first wave meiosis initiation and *Foxl2* positive pregranulosa cells may form follicles that grow directly after birth (first wave of folliculogenesis), while the cortical germ cells and *Lgr5* positive pregranulosa cells that subsequently express *Foxl2* may form dormant primordial follicles that need to be activated before growth (second wave of folliculogenesis).

### Regional characters of granulosa cells in the postnatal mouse ovary

We next turned our attention to the postnatal follicle development, and the spatial distribution of *Ooep* (germ cells), *Inha*, *Inhbb* and *Fst* (granulosa cells) and *Star* and *Cyp17a1* (theca cells) were examined by BayesSpace at PD7, PD14, 1M and 2M stages (Fig. 5A and fig. S5A). Notably, *Cyp17a1* is absent in the ovary at the PD7 stage. Additionally, Inha is widely expressed in the ovary and exhibits a higher expression level in large follicles (Fig. 5A and B). Furthermore, the *Inhbb* and *Fst* are highly expressed in large follicles (Fig. 5A). To further dissect the molecular heterogeneity of granulosa cells during follicle growth, we re-clustered the granulosa cells of PD90 into four groups; we annotated them as progenitor granulosa cells (pGC, *Kctd14*), proliferative granulosa cells (ProGC, *Top2a*), cumulus granulosa cells (cGC, *Inhbb*) and mural granulosa cells (mGC, *Prss23*) (Fig. 5C and D). To gain further insight into the transcriptome differences among the subclusters of granulosa cells, we compared the top 200 expressed DEGs among the four subtypes of granulosa cells. The Venn diagram demonstrated that there were no overlapped DEGs among the four subtypes of granulosa cells, and different two subclusters shared a very low percentage of overlapped DEGs (fig. S5B). Moreover, GO analysis of the nonoverlapped DEGs (145, 182, 118 and 156 in fig. S5B) was performed, revealing that the GO terms “alpha-actinin binding” and “hormone binding” were specific to the cGC cluster, while “cell adhesion molecule binding” and “NAD binding” were specific to the mGC cluster (fig. S5C).

**Figure 5.**
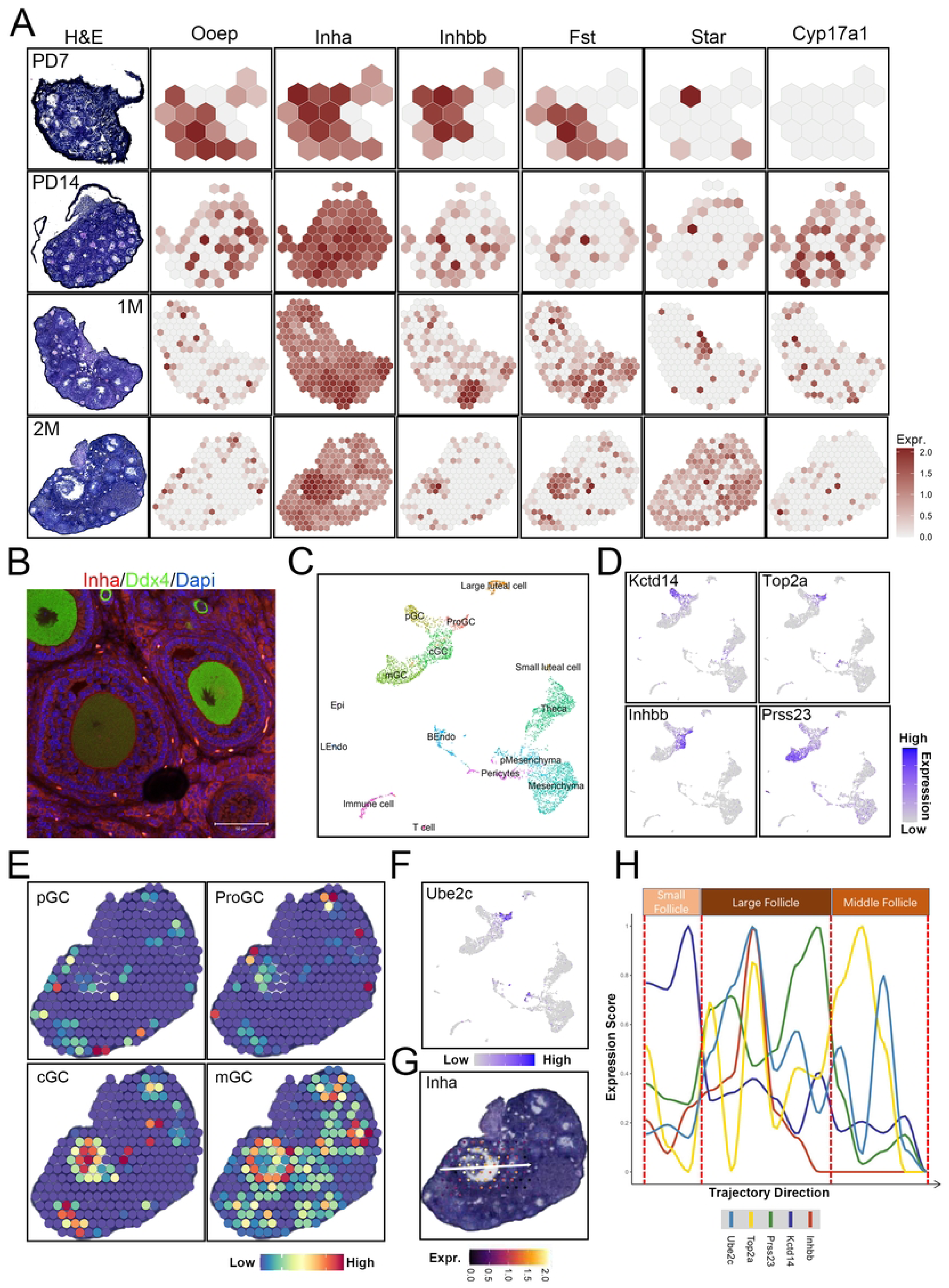
Spatial transcriptomics identify spatially restricted genes during follicle development. (A) Spatially resolved gene expression for follicle development. H&E, hematoxylin and eosin. (B) Immunostaining for Inha and Ddx4 in mouse ovary. (C) Visualization of major clusters of PD90 ovarian cells using UMAP. BEndo, blood associated endothelial cells; cGC, cumulus granulosa cells; Epi, epithelial cells; Lendo, lymph associated endothelial cells; mGC, mural granulosa cells; pGC, progenitor granulosa cells; pMesenchyma, proliferative mesenchyma; ProGC, proliferative granulosa cells. (D) UMAP showing expression patterns of selected markers for subclusters of granulosa cells in Figure 5C. Purple (or gray) represents a high (or low) expression level as shown on the color key at the right bottom. (E) Visualization of granulosa cell subclusters on 2-month mouse ovary based on integration of scRNA-seq cell type annotations with spatial transcriptomic data. (F) The expression patterns of Ube2c in PD90 ovarian cells. (G) The expression patterns of Inha along trajectory. (H) Dynamic expression patterns of follicle development associated genes along trajectory.

To further dissect the spatial distribution of different subtypes of granulosa cells, we mapped these four subclusters of granulosa cells to the 2M ovary section and found that the pGC mainly located at the ovary cortex, the cGC mainly located at the cumulus-oocyte complex and the mGC specifically highly located at the peripheral regions of mature follicles (Fig. 5E). Interestingly, a subset of ProGC overlapped with pGC and the other ProGC overlapped with cGC, suggesting that both pGC and cGC are highly proliferative (Fig. 5E and fig. S5D). Moreover, *Ube2c* is highly expressed in the ProGC (Fig. 5F). We next sought to understand the unique features of granulosa cells in different sizes of follicles. The spatial trajectory was assigned using SPATA (Fig. 5G), and the expression patterns of known marker genes for different subtypes of granulosa cells were examined along the trajectory. In small follicles, *Kctd14* is highly expressed and then downregulated in large and middle follicles. Additionally, the *Inhbb*, *Top2a* and *Ube2c* are consistently highly expressed in large follicles (mature follicles), while *Inhbb* is significantly downregulated in middle follicle in spite of highly expressed *Top2a* and *Ube2c*, suggesting that *Inhbb* may play an important role in promoting follicle maturation (Fig. 5H). To further understand the molecular differences between subordinate and dominant follicles, *Grb14*, a marker gene for subordinate follicles [30], was examined in 2M, 1M and PD14 ovary sections (fig. S5E), which revealed that *Grb14* was highly expressed in the subordinate follicle in the 1M ovary section (fig. S5E), while in the contiguous dominant follicle the *Grb14* is significantly downregulated (fig. S5E). To confirm this result, we constructed a spatial trajectory throughout the subordinate and dominant follicles in 1M ovary section, and the expression patterns of relevant genes were examined (fig. S5F). We revealed that *Grb14* was highly expressed in subordinate follicles and significantly downregulated in dominant follicles, while *Inhbb* was lowly expressed in subordinate follicles and significantly upregulated in dominant follicles, suggesting that *Grb14* and *Inhbb* play important roles in follicular deviation (fig. S5F). Above all, through the single-cell and spatial transcriptomes, we precisely identified several categories of granulosa cells with region-specific properties, which are closely associated with the various physiological functions of follicle development. Our findings highlight the molecular differences between subordinate and dominant follicles from a novel developmental perspective.

### Spatial characteristics of the corpus luteum-specific luteal subtypes

We next turned our attention to the region-specific luteal cells, and two common cell clusters of the corpus luteum, LLC and SLC, have been identified (Fig. 5C). To comprehensively characterize the molecular signatures between LLC and SLC, the highly DEGs were examined; *Ssu2* and *Bhmt* expressions were specifically enriched in LLC, while *Onecut2* and *Akr1c18* expressions were specifically enriched in SLC (Fig. 6A). Moreover, *Ptgfr* and *Sfrp4* were highly expressed in both LLC and SLC (Fig. 6A). Furthermore, with the aid of structural ovary markers, spatial transcriptomic imaging located the LLC both in the corpus luteum and other ovary regions, while the SLC specifically located in the corpus luteum (Fig. 6B). To further exclude the impacts of potential precursor cells, other cell clusters were also mapped to the spatial ovary sections, and all of them exhibited none or low enrichment in the corpus luteum (Fig. 5E and fig. S6A and B). To interpret the functional differences, we compared the top 200 expressed DEGs between LLC and SLC. The Venn diagram showed that there were 51 shared DEGs between LLC and SLC (fig. S6C). Moreover, GO analysis of the nonoverlapped and shared DEGs was performed; the GO terms “protein kinase inhibitor activity” and “hormone binding” were specific to SLC, “cytochrome-c oxidase activity” and “steroid dehydrogenase activity” were specific to LLC, and “peroxidase activity” and “antioxidant activity” were highly enriched in shared DEGs between SLC and LLC (fig. S6D).

**Figure 6.**
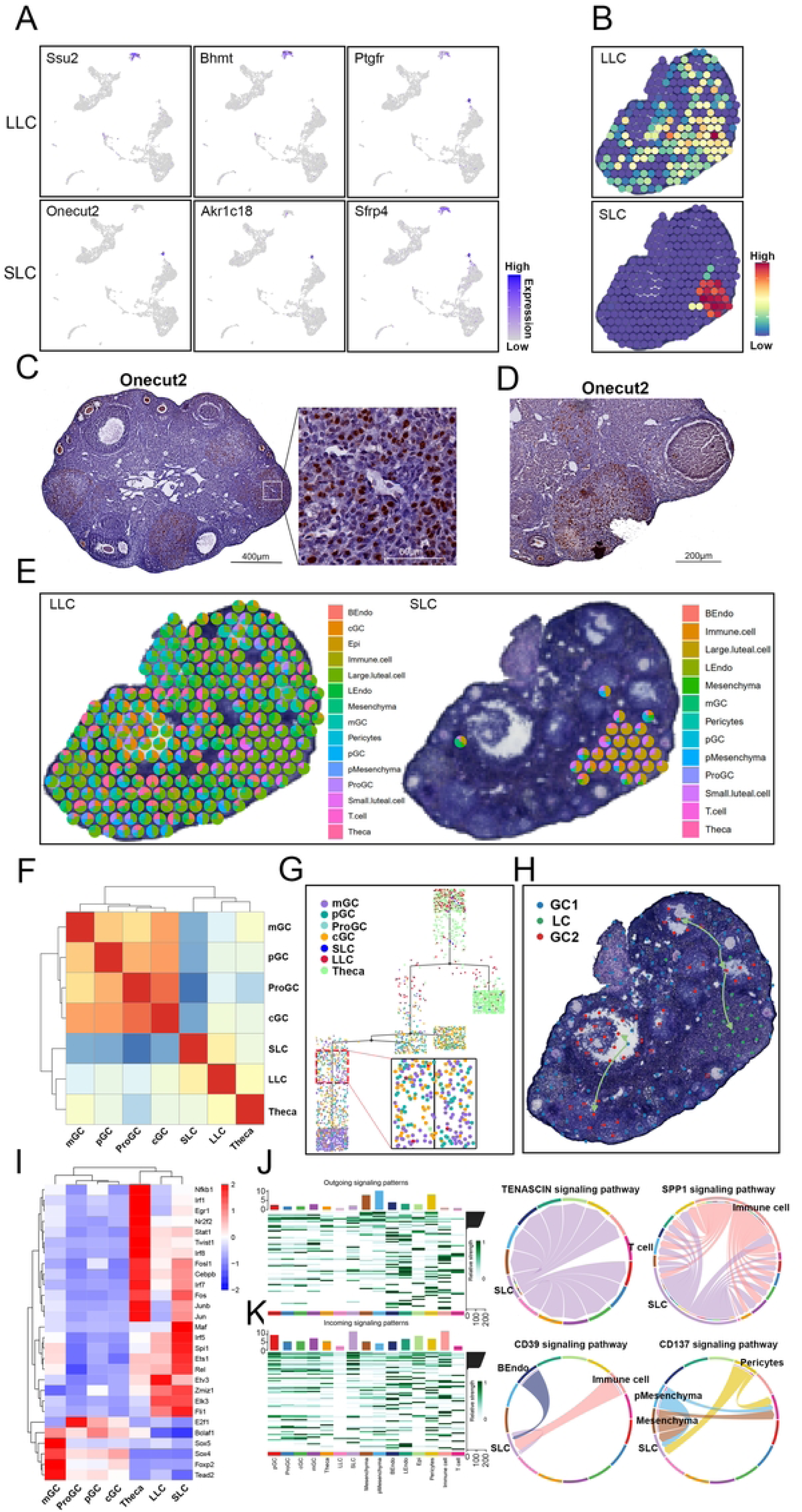
Molecular diversity and spatial distribution differences of luteal cells in mouse ovary. (A) UMAP visualization of marker genes for LLC and SLC. Color key represents the expression levels. LLC, large luteal cells; SLC, small luteal cells. (B) Visualization of LLC and SLC on 2-month mouse ovary based on integration of scRNA-seq cell type annotations with spatial transcriptomic data. (C) Immunohistochemistry staining of Onecut2 in mouse ovary. The scale bars represent 400 μm in low-magnification view and 60 μm in high-magnification view. (D) Immunohistochemistry staining of Onecut2 during ovulation. The scale bars represent 200 μm. (E) Spatial scatter pie plot representing the proportions of the LLC (left) and SLC (right) from the PD90 ovary reference atlas in the 2-month mouse ovary. (F) Heatmap showing the correlations among different cell clusters by Spearman analysis. (G) Pseudotime trajectory of granulosa cells, luteal cells and Theca cells analyzed by Monocle. (H) Spatial trajectory showing the differentiation of granulosa cells analyzed by stLearn. (I) Heatmap of regulon activity analyzed by SCENIC. (J) The outgoing signaling patterns among different cell clusters (left); The typical outgoing signaling pathway of SLC (right). (K) The incoming signaling patterns among different cell clusters (left); The typical incoming signaling pathway of SLC (right).

Based on the specific enrichment of *Onecut2* expression in SLC, to experimentally validate its expression pattern, we performed immunohistochemistry using antibodies against Onecut2, which showed that *Onecut2* was highly expressed in SLC (Fig. 6C). Surprisingly, we also found that *Onecut2* was particularly expressed in the oocytes during follicle development, which was further verified by western blot results (Fig. 6C and fig. S6E). To further detect the expression time of *Onecut2* in the corpus luteum, we also examined the expression patterns of *Onecut2* in a series of ovary sections, which showed that the corpus luteum begins to express *Onecut2* as ovulation occurs (Fig. 6D). To deconvolute the spatial spots located in the corpus luteum, we integrated the single-cell and spatial transcriptomic data and inferred the ratios of cell types within the spots using SPOTlight [31]. We revealed that the majority of cells in the corpus luteum are LLC, and approximately 35% cells are the SLC (Fig. 6E and fig. S6F). Moreover, spatial interactions among different cell types also indicated that LLC is close to the theca, mGC and pericytes, respectively (fig. S6G).

To further understand the relationships between potential precursor cells and the LLC and SLC, we performed correlation analysis among the subtypes of granulosa cells, theca cells, LLC and SLC, which showed that the four subtypes of granulosa cells were clustered together, while the theca cells, LLC and SLC were clustered together (Fig. 6F). Moreover, the SLC were significantly uncorrelated to granulosa cells, suggesting that the granulosa cells may not be the precursor cells of SLC (Fig. 6F). To explore the derivation of LLC and SLC, we also constructed a pseudotime trajectory using monocle, which revealed both granulosa cells and theca cells contain LLC, but only theca cells contain SLC, suggesting that SLC may derive from theca cells, while LLC may derive from granulosa cells and theca cells (Fig. 6G). Moreover, we also constructed pseudo-space-time trajectories using stLearn [23], which showed a well differentiation process from granulosa cells to luteal cells (Fig. 6H and fig. S6H). Subsequently, we examined the spatial distribution of typical transition marker genes, such as *Adgrg1*, *Slc38a5* and *Rab27a*, which showed a higher expression in specific regions (Fig. 6I). To further explore the dynamics of transition marker genes, we constructed a spatial trajectory using SPATA, which showed that *Rarres2* and *Scaf1* were highly expressed in small follicles, *Ctgf* and *Scl38a5* were highly expressed in middle follicles and *Rab27a*, *Nrcam* and *Foxk2* were highly expressed in the corpus luteum, respectively (fig. 6I and J).

Next, we performed SCENIC analysis and discovered dozens of transcription factors with a different enrichment pattern. Hierarchical clustering analysis revealed the regulatory relationship between granulosa cells and luteal cells (Fig. 6I). For instance, Sox4 and Foxp2 were active mainly in the granulosa cells, while Junb and Cebpb were active mainly in the luteal cells and theca cells. Moreover, Etv3 are highly enriched in the LLC and Maf are highly enriched in the SLC (Fig. 6I). To further dissect the roles of SLC, we then performed CellChat analysis to investigate the cell interactions between SLC and other cell clusters. Among the outgoing signaling patterns, TENASCIN signaling pathway is highly enriched in the SLC, which participate in the regulation of other cell clusters except T cells (Fig. 6J). Notably, SLC is the top cluster for incoming signaling patterns, and Bendo, immune cell and mesenchyma could regulate physiological function of SLC through CD39 and CD137 signaling pathway (Fig. 6K). Collectively, we identified a novel corpus luteum specific cell type (SLC) that may derive from theca cells, and we compared the molecular and functional differences between LLC and SLC, which provided important new information for understanding the physiological function of the corpus luteum.

## Discussion

In this study, we systematically analyzed the single-cell and spatial transcriptomic maps of the mouse ovary from the early fetal gonad to the adult mature ovary to understand the cellular and molecular diversity during ovarian cell lineages differentiation and follicle formation and development. Overall, we constructed pseudotime trajectories to show the two waves of oogenesis and cell fate divergency during BPG differentiation. By combining spatial and scRNA-seq assays using bioinformatic approaches, we explored the relationship between two waves of meiosis initiation and folliculogenesis. Moreover, we also dissected the molecular signatures and spatial distribution of granulosa cell subtypes to understand the processes of follicular growth and deviation. Strikingly, we discovered a new cell type (SLC) that highly expressed *Onecut2* and specifically located in the corpus luteum. Altogether, we here present the single-cell and spatial transcriptomic atlas of mouse ovaries from early fetal to adult stages, which provides mechanistic insights into both molecular and morphological aspects of ovary development.

Early female germ cell development mainly involves meiosis initiation and arrest, which exhibits two waves of oogenesis. Our previous study has revealed two waves of oogenesis in monkey fetal ovaries, and the germ cells undergoing death around the second wave of oogenesis significantly expressed *Spo11* and *Prdm9* [11]. In this study, we constructed the pseudotime trajectory in mouse female germ cells, which also exhibited two waves of oogenesis, suggesting that the basic events during oogenesis are conserved between mouse and monkey (Fig. 2C). Moreover, the germ cells undergoing death around the second wave of oogenesis in the mouse also highly expressed *Prdm9* consistent with that in monkey germ cells (Fig. 2D). Furthermore, the dying germ cells also specifically expressed *Pbx3*, a transcription factor, suggesting an important role of *Pbx3* in the regulation of oocyte attrition (fig. S2D). Additionally, the proper differentiation of ovarian somatic cells is essential for germ cell development [32]. Therefore, we also systematically analyzed the differentiation of BPG that further gave rise to theca GC in branch 1, which is consistent to the lineage tracing results in *Foxl2* positive cells [33]. Notably, the facet plots of FGC and BPG trajectories showed that the time for the BPG giving rise to theca GC is similar to that of the second wave of oogenesis (Fig. 2E and Fig. 3F), suggesting a close relationship between theca cell differentiation and oocyte attrition or survival.

Meiosis initiation is a critical event during gametogenesis. A previous study has reported that retinoic acid is required for *Stra8* expression which could promote the mitosis to meiosis transition [34]. However, a recent study found that meiosis occurs normally in the fetal ovary of mice lacking all retinoic acid receptors [35]. In addition, transcriptional factors, such as *Msx1*, *Msx2* and *Zglp1* could promote meiosis initiation through activating the downstream effector Stra8 [27, 36]. Interestingly, gonad somatic cells have been shown to be involved in germ cell meiosis initiation, and abnormal differentiation of gonad somatic cells could cause meiosis initiation defects [32]. Our study provides insights into the meiosis initiation of female germ cells. The data suggest that female germ cells initiate meiosis first in the medullary regions and then in the cortical regions of the ovary (Fig. 4A), which is consistent with recent studies [25, 37]. Moreover, *Dazl* is highly expressed in the medullary regions where *Stra8* is also highly expressed but *Dppa5a* expression is downregulated (Fig. 4A), which has been shown by our previous study [38]. What triggers the female germ cells to switch from mitosis to meiosis at specific developmental times and locations remains to be fully elucidated but our data provides a dynamic atlas for meiosis initiation at single-cell and spatial transcriptomic levels.

Single-cell RNA sequencing is a powerful tool for the systematic identification of cell types in a given tissue but it does not preserve its spatial information, leading to an inaccurate characterization of cell types and predictions of their physiological function. Our analysis demonstrates that spatial transcriptomics can be used to improve the robustness of scRNA-seq. In this study, we accurately characterized the subtypes of granulosa cells, mesenchyma and luteal cells (Fig. 5E and Fig. 6B and fig. S6A and B), considering their known marker genes and spatial distributions. For instance, mesenchyma mainly located in the mesovarium, while proliferative mesenchyma (pMesenchyma) mainly located in the ovarian parenchyma (fig. S6A), suggesting an important role for pMesenchyma in the reshaping of the ovarian microenvironment as waves of follicular growth.

The important physiological function of the corpus luteum is producing the hormone progesterone to support pregnancy, but the cell compositions of the corpus luteum and their spatial distributions are still poorly understood. Our research dissected the subtypes of luteal cells and identified a novel cell type (SLC) that highly expressed *Onecut2* and specifically located in the corpus luteum (Fig. 6A and B). Interestingly, immunostaining showed that Onecut2 is specifically expressed in the oocyte during follicular growth and then specifically expressed in the SLC as ovulation occurs (Fig. 6C and D). A previous study demonstrated that Onecut2 could suppress the androgen receptor transcriptional program by direct regulation of androgen receptor target genes [39]. Therefore, the transition of *Onecut2* expression from the oocyte to SLC and the functions of Onecut2 in the corpus luteum need to be further explored.

In conclusion, we have combined single-cell and spatial transcriptomics to explore the morphogenesis of the mouse ovary at high molecular, spatial, and temporal resolution. We analyzed cellular proximities within the ovary tissue, measured changes in cellular composition with anatomical location, and quantified anatomically restricted gene expression. Because cellular transitions in complex lineages do not occur in a synchronized manner, our data represent a broad range of cellular states from the undifferentiated gonad to the adult ovary. We mapped information of cell lineage differentiation transitions that were obtained from pseudotime ordering of single-cell transcriptomes, on spatial ovary sections, and we were thereby able to demonstrate how cellular differentiation and morphological changes co-occur. Our research thus demonstrates how combined spatial and single-cell RNA sequencing can be used to study the interplay between cellular differentiation and morphogenesis.

## Materials and Methods

### Ethics statement

WT C57BL/6J mice were purchased from Beijing SPF Biotechnology Co., Ltd. All mice were housed in a specific pathogen-free facility with individually ventilated cages and free access to a regular rodent chow diet and water. The room had a controlled temperature (23~25 °C), controlled humidity (40~65%) and light program (alternating light/dark cycles). All experiments and methods in this study were kept in compliance with the guidelines of the Animal Care and Use Committee of the Institute of Zoology at the Chinese Academy of Sciences. Gestational age of embryos was determined by checking vaginal plugs, with noon of the day of the plug appearance defined as E0.5.

### Preparation of ovarian cell suspensions and cryosections

The ovary samples for scRNA-seq were dissected from female mice at PD0, PD21 and PD90 days. For each single-cell sequencing experiment, the clean ovaries were subjected to a standard digestion procedure through Tumor Dissociation Kit (MiltenyiBiotec #130-095-929) as described previously [11]. Finally, the ovarian cell suspensions were prepared for further library construction. For the 10× Visium experiment, the fetal and postnatal ovaries were collected from female C57BL/6J mice at E13.5, E18.5, PD0, PD3, PD7, PD14, PD30 and PD60. The ovary samples were embedded with pre-chilled optimal cutting compound (OCT) and the frozen ovaries were sectioned at 10 µm thickness. Subsequently, each ovarian cryosection was mounted on 10x Visium Spatial Tissue Optimization slides.

### Single-cell and spatial transcriptomic libraries construction and sequencing

The suspended single ovarian cells were encapsulated into droplet emulsions using a 10× Chromium system. Then, the processes of library construction were instructed by the manufacturer’s protocol of Chromium Single-Cell 3′ Gel Bead and Library V3 kit. After the generation of GEMs, reverse transcription reactions were barcoded using a unique molecular identifier (UMI), and 12 cycles were used for cDNA amplification. The resulting libraries were fragmented and assessed on a fragment analyzer using a High Sensitivity NGS Analysis Kit. The average fragment length was quantified using fragment analyzer and qPCR with a Kapa Library Quantification Kit for Illumina. The final libraries were sequenced on an Illumina NovaSeq6000 (Illumina, San Diego) with pair-end 150 bp (PE150) reads.

For 10X Visium libraries construction, the sections were first placed on Thermocycler Adaptor with the active surface facing up and incubate 1 min at 37°C, and then fixed for 30 min with methyl alcohol in −20°C followed by staining with hematoxylin and eosin. The brightfield images were taken on a Leica DMI8 whole-slide scanner at 10X resolution. Next, permeabilization was performed using permeabilization enzyme to release spot cellular mRNA followed by reverse transcription for cDNA synthesis. Subsequently, the cDNA amplification was performed on a S1000TM Touch Thermal Cycler (Bio Rad). Finally, the Visum spatial libraries were constructed using Visum spatial Library construction kit (10x Genomics, PN-1000184), which were further sequenced on an Illumina Novaseq6000 with a sequencing depth of at least 100,000 reads per spot.

### Processing of single cell RNA-seq data

Cell Ranger v4.0.0 was used to perform raw data demultiplexing, reads mapping and barcode processing to generate a matrix of gene counts versus cells. Briefly, the raw BCL files generated by Illumina NovaSeq 6000 sequencer were demultiplexed into fastq files by function mkfastq. The fastq files were then processed using the count function with default settings, including alignment (using STAR align to mm10 mouse reference genomes), filtering, and UMI counting. The generated count matrices were loaded into R using the Read10X function of Seurat package (version 4.1.0)[40]. The Seurat object was created based on two filtering parameters of “min.cells = 5” and “low.thresholds = 200”, followed by adding the sample information in the row names of the matrices and integrating data from different samples. Next, the Seurat object was processed with harmony to remove batch effects [21]. Subsequently, normalization was performed in Seurat on the filtered matrix to obtain the normalized count and principal component analysis (PCA) was performed to reduce the dimensionality on the top 18 principal components. Then, cells were clustered at a proper resolution and visualized in two-dimensions using UMAP [41]. To identify typical marker genes for cluster annotation, the “FindAllMarkers” function was applied to detect differentially expressed genes between clusters with default parameters.

### Processing of 10×Visium data

Space Ranger v1.2.0 was used to demultiplex Visium-prepared raw data by function mkfastq. The output fastq files and microscope slide images were then run with function count using default settings, including alignment (using STAR align to mm10 mouse reference genomes), tissue detection, fiducial detection, and barcode counting. The generated feature-barcode matrices were loaded into R using the Load10X_Spatial function of Seurat. According to the developer’s vignettes, the data were filtered and normalized by SCTransform [42]. Spatially variable genes were identified using the method of markvariogram and PCA was performed to reduce the dimensionality on the top 30 principal components. Spots were clustered based on the stSME clustering by stLearn [23]. Additionally, the SPOTlight package was used to deconvolute spatial transcriptomics spots and infer the ratio of cell types within a certain spot [31].

### Pseudotime and Pseudo-space-time analysis

Pseudotime analysis was performed on germ cells and granulosa cells from the mouse ovary atlas data by using the Monocle2 R package [43]. With the gene count matrix as input, the new dataset for Monocle object was created. The ordering genes were differentially expressed genes between clusters in each cell type, which was further used to recover lineage trajectories. After pseudotime time was determined, differentially expressed genes were clustered to verify the fidelity of lineage trajectories. The root state was set and adjusted following consideration of the biological meanings of different cell branches. Additionally, pseudo-space-time analysis was performed on 2M ovary sections by using stLearn [23]. The ordering genes were differentially expressed genes between clades, which was further used to construct spatial trajectories.

### Cell-cell interaction analysis

CellChat objects were created via Cellchat (https://github.com/sqjin/CellChat, R package, v.1). With ‘‘CellChatDB.mouse’’ set up as the ligand-receptor interaction database, cell-cell interaction analysis was then performed via the default setting.

### GO analysis

GO enrichment analyses were performed using ClusterProfiler, an R package in Bioconductor, to detect the gene-related biological process, and GO terms with a threshold value of “pvalue Cut off = 0.05” were considered.

### Transcription factor regulatory network analysis

SCENIC v1.3.1 was used to predict the core regulatory transcription factors and their target genes based on the scRNA-seq data[44]. Following the standard pipeline, the co-expressed genes for each transcription factor were identified via the GENIE3 (v1.18.0) [45]. RcisTarget was used to infer enriched transcription factor-binding motifs and to predict candidate target genes (regulons) based on the mm10 mouse-specific database containing motifs with genome-wide rankings. Finally, regulon activity was calculated by AUCell software.

### Immunofluorescence analysis

For immunostaining, the paraffin sections were dewaxed and rehydrated in xylene and in a series of decreasing graded ethanol to PBS. For antigen retrieval, the sections were treated in citric acid for 20 min at 98°C. After cooling down, the sections were incubated with 5% BSA in 0.3% Triton X-100 for 1 hour and were then incubated overnight at 4°C with the primary antibodies. The following primary antibodies were used: Cdx2 (Abcam, ab76541), Inha (Invitrogen, MA5-15703), Ddx4 (Abcam, ab13840), and Ddx4 (Abcam, ab27591). After being washed three times in PBS, the sections were incubated with secondary antibodies for 1 hour at room temperature. Nuclei were stained with 4′,6-diamidino-2-phenyl-indole (DAPI, Life Technologies) and sections were examined with a confocal laser scanning microscope (Carl Zeiss Inc, Thornwood, NY).

### Immunohistochemistry analysis

For paraffin-embedded ovarian tissues, the sections were dewaxed and rehydrated in xylene and in a series of decreasing graded ethanol to PBS. Next, the sections were pretreated in citric acid for 20 min at 98°C and cooled down to room temperature and then were incubated in 0.3% H_2_O_2_ for 10 min to block endogenous peroxidase activity. The sections were incubated overnight at 4°C with primary antibody Onecut2 (Proteintech, 21916-1-AP) and 1% PBS was used to wash the sections three times for 5 min each on the next day. Then the sections were incubated with secondary antibodies for 20 min at room temperature. The sections were washed by 1%PBS three times for 5 min each. Then the DAB was added to each section by incubating for 1-2 min and hematoxylin was used for counterstaining for 1-2 min. Then the sections were dehydrated through a series of increasing graded ethanol, cleared in xylene, and sealed. Brown staining of the cytoplasm or nucleus of the cells was considered as positive.

### Western blot analysis

The ovary and liver tissues were washed with PBS and lysed with RIPA buffer supplemented with protease inhibitors cocktail (Roche). Equal amounts of total protein were boiled in sodium dodecyl sulfate (SDS) sample buffer for 5 min. The boiled proteins were separated by SDS/PAGE gels and then transferred to PVDF membranes. The blots were probed with the primary antibody at an appropriate dilution by overnight incubation at 4 °C, followed by 1-h incubation with appropriate HRP-conjugated secondary antibodies at room temperature. The protein expression was normalized to that of Gapdh. The antibodies used were Onecut2 (Proteintech, 21916-1-AP).

### Data availability

The raw sequence data reported in this paper have been deposited in the Genome Sequence Archive in National Genomics Data Center, China National Center for Bioinformation/Beijing Institute of Genomics, Chinese Academy of Sciences (GSA: CRA010027) that are publicly accessible at https://ngdc.cncb.ac.cn/gsa.

## ACKNOWLEDGMENTS

This work was supported by National R&D program of China, Grant/Award Number:2022YFC2703501; Natural Science Foundation of Shandong Province, Grant/Award Number ZR2021ZD33; and Science and Technology Program of Guangzhou, China, Grant/Award Number 202201020292.

## AUTHOR CONTRIBUTIONS

Zheng-Hui Zhao and Qing-Yuan Sun conceived and supervised the project, designed the experiments and wrote our manuscript. Tie-Gang Meng collected and processed ovary samples. Zheng-Hui Zhao conducted computational analysis and validation experiments. Fei Gao provided technical assistance and Heide Schatten polished the language of our manuscript.

---

**Supplementary Figure S1.**
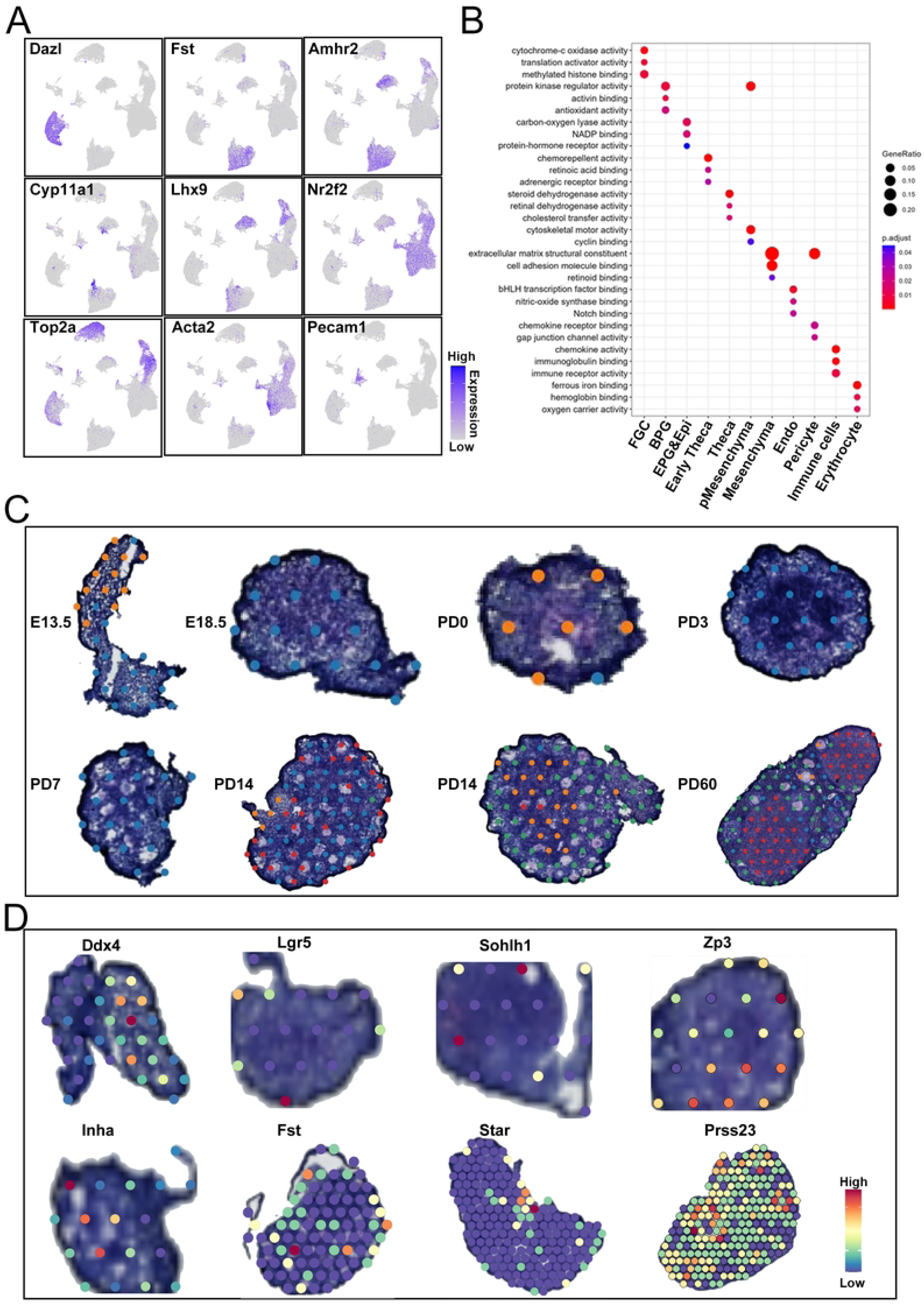
Single-cell and spatial transcriptomics identify distinct cell clusters. (A) UMAP visualization of marker genes for ovarian cell clusters. A gradient of gray to purple indicates the low to high gene expression level. (B) Diagram showing the enrichment of GO terms in eleven ovarian cell clusters. The dot size is positively correlated with the gene ratios. (C) Spatially resolved gene expression for region restricted genes. (D) Spatial RNA-seq spots clustered by stLearn.

**Supplementary Figure S2.**
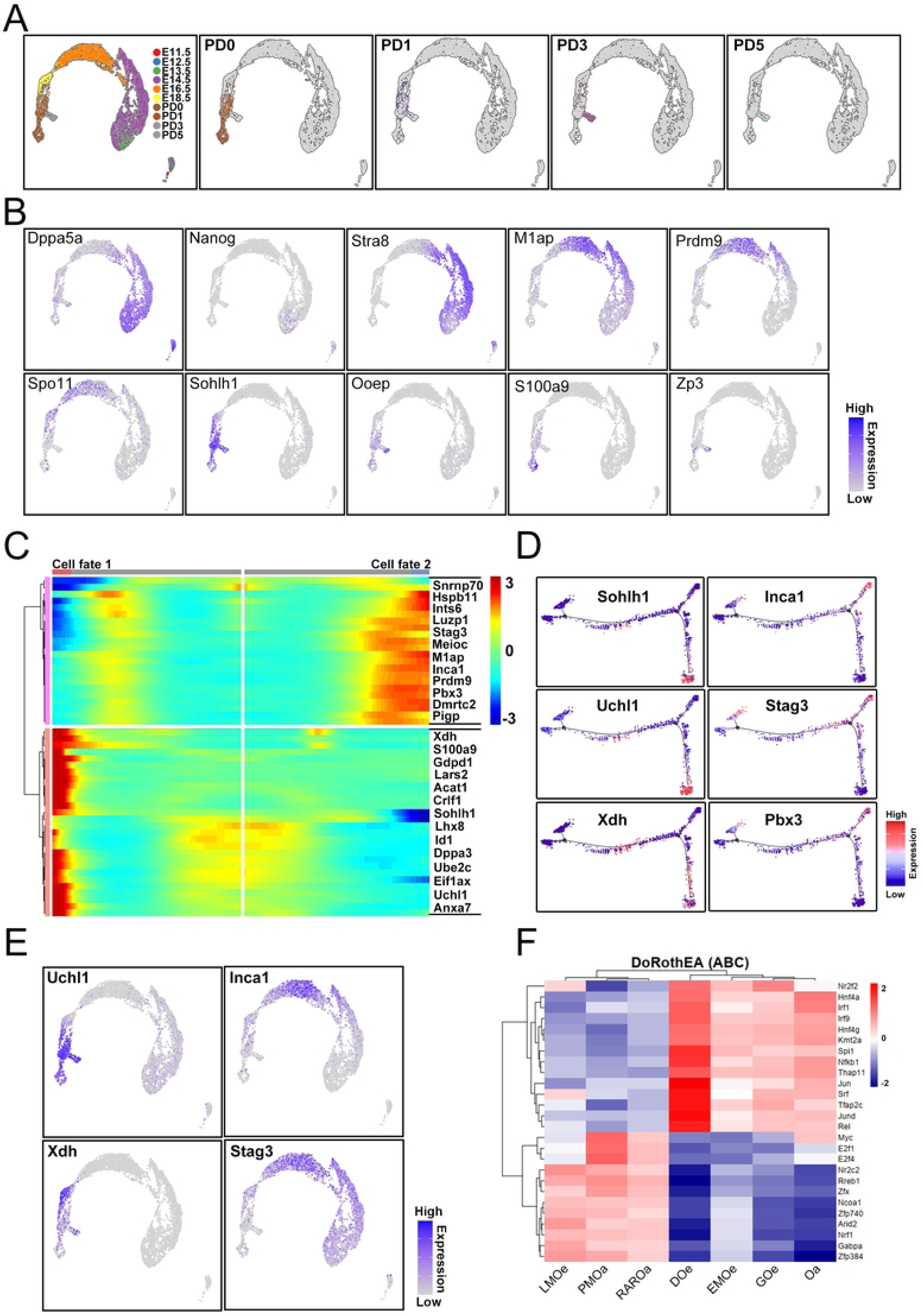
Molecular signatures of female germ cells during early oogenesis. (A) UMAP projection of female germ cells colored by developmental stages. (B) Expression patterns of selected markers for female germ cell subclusters. Purple (or gray) represents a high (or low) expression level as shown on the color key at the right bottom. (C) Heatmap representing the expression dynamics of 2 gene sets with increased or reduced expression at the cell fate 1 (oocyte survival) and cell fate 2 (oocyte attrition) stages. (D) Expression of representative genes in each set along single-cell pseudotime trajectories. (E) UMAP visualization of representative genes in each set. A gradient of gray to purple indicates the low to high gene expression level. (F) Heatmap of highly variable transcription factor activities among the seven groups. Doe, dormant oocyte; EMOe, early meiotic oocyte; GOe, growing oocyte; LMOe, late meiotic oocyte; Oa, oogonia; PMOa, pre-meiotic oogonia; RAROa, RA responsive oogonia.

**Supplementary Figure S3.**
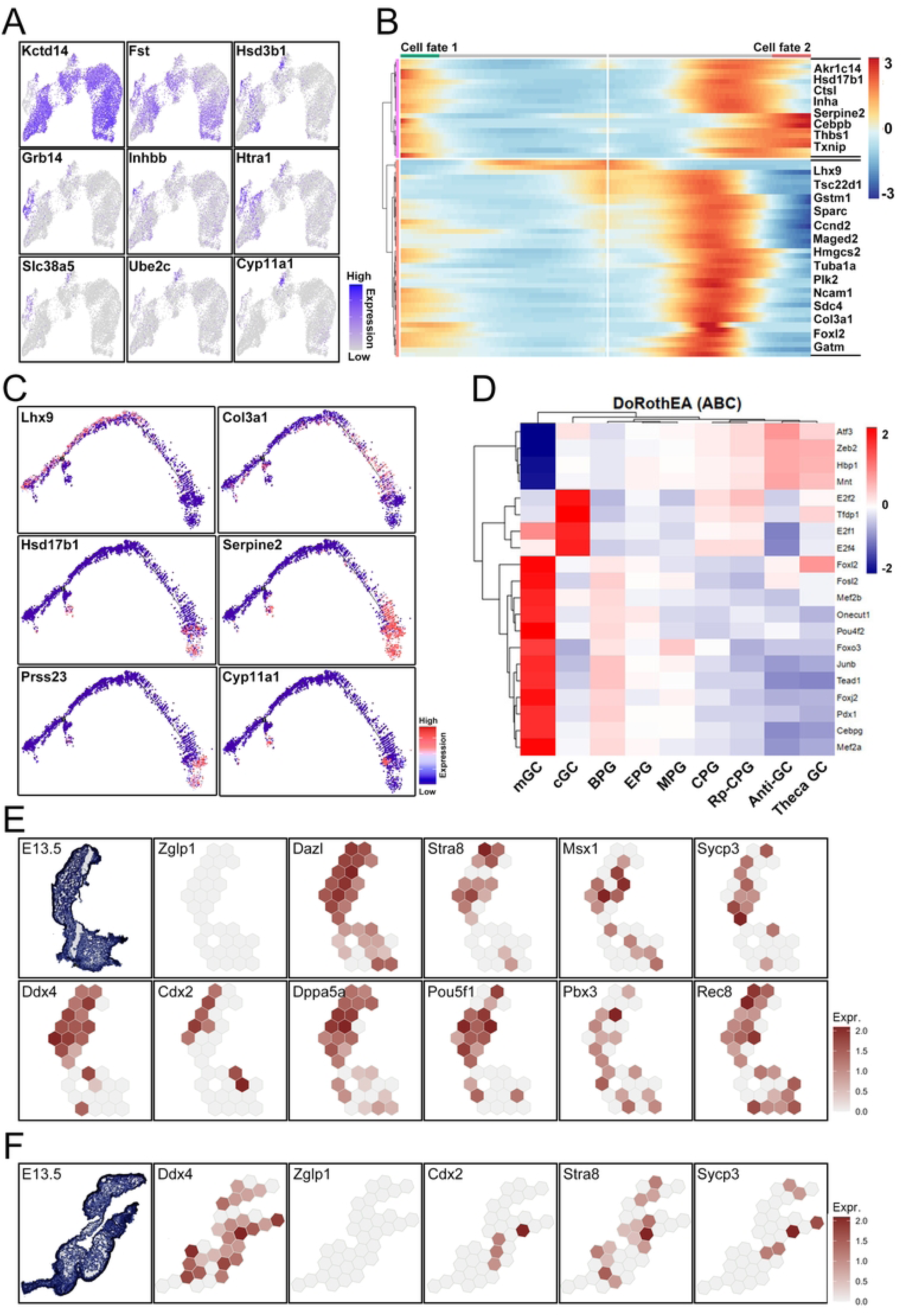
Molecular diversity of granulosa cells during differentiation processes. (A) Expression patterns of selected markers for granulosa cell subclusters. Purple (or gray) represents a high (or low) expression level as shown on the color key at the right bottom. (B) Heatmap representing the expression dynamics of 2 gene sets with increased or reduced expression at the cell fate 1 (Theca cells) and cell fate 2 (granulosa cells) stages. (C) Expression of representative genes in each set along single-cell pseudotime trajectories. (D) Heatmap of highly variable transcription factor activities among the nine groups. Anti-GC, antioxidative granulosa cells; BPG, bipotential pregranulosa cells; cGC, cumulus granulosa cells; CPG, cortical pregranulosa cells; EPG, epithelial pregranulosa cells; MPG, medullary pregranulosa cells; mGC, mural granulosa cells; Rp-CPG, ribosomal protein genes associated cortical pregranulosa cells; Theca GC, granulosa cells that differentiated into Theca cells. (E) Spatially resolved gene expression for transitions from mitosis to meiosis associated genes enhanced by BayesSpace. (F) Spatially resolved expression patterns of meiosis associated genes enhanced by BayesSpace.

**Supplementary Figure S4.**
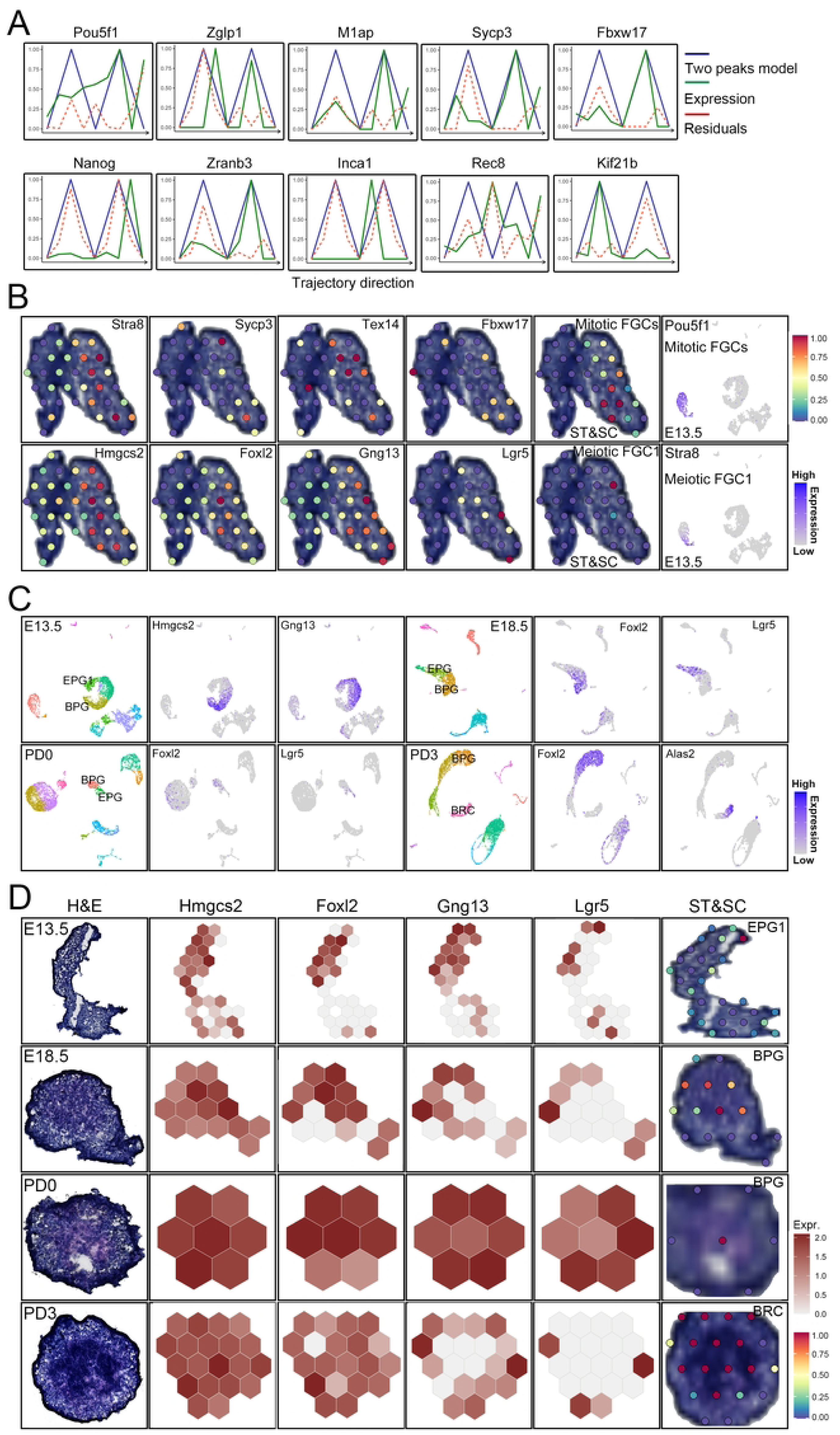
Characteristics of different spatial regions during early ovary development. (A) Visualization of differentially expressed genes along the trajectory in Figure 4C. (B) Spatially resolved gene expression for region restricted genes (left); Visualization of mitotic FGCs and meiotic FGC1 on E13.5 mouse ovary based on integration of scRNA-seq cell type annotations with spatial transcriptomic data (right). (C) Visualization of major clusters of E13.5, E18.5, PD0 and PD93 ovarian cells and associated marker genes using UMAP. A gradient of gray to purple indicates the low to high gene expression level. BPG, bipotential pregranulosa cells; EPG, epithelial pregranulosa cells; BRC, blood related cells. (D) Spatially resolved gene expression patterns for cortical and medullary follicle formation. H&E, hematoxylin and eosin; ST&SC, spatial transcriptomics and single cell transcriptomes.

**Supplementary Figure S5.**
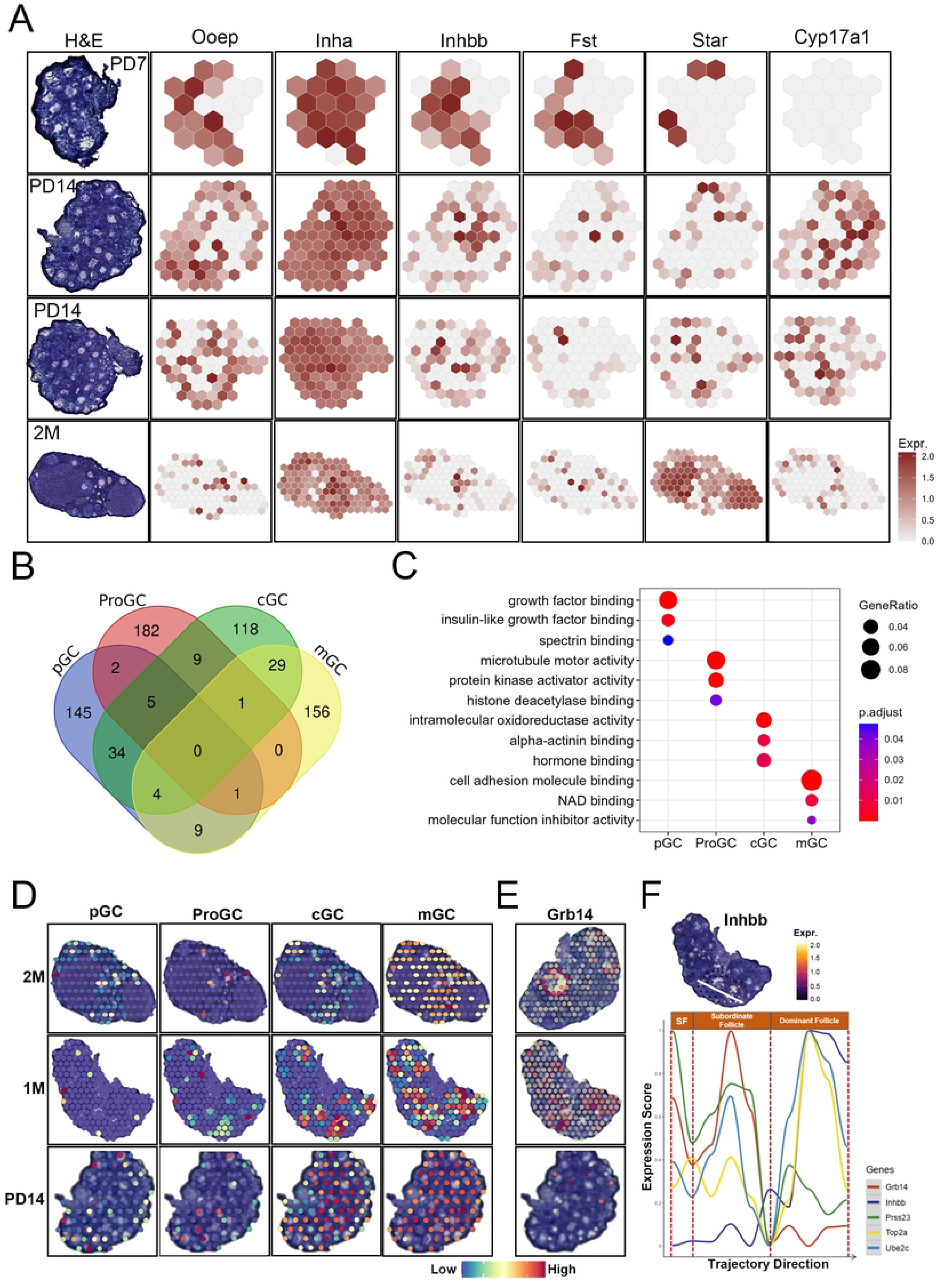
Spatially resolved gene expression for follicle growing. (A) Spatially resolved gene expression for follicle growing. H&E, hematoxylin and eosin. (B) Venn diagram of top 200 differentially expressed genes in pGC, ProGC, cGC and mGC demonstrates overlap among the gene populations in four subclusters of granulosa cells. (C) The top GO terms enriched in the differentially expressed genes that unique to pGC (145), ProGC (182), cGC (118) and mGC (156) in Figure S5B. (D) Visualization of pGC, ProGC, cGC and mGC on PD14, 1-month and 2-month mouse ovary respectively based on integration of scRNA-seq cell type annotations with spatial transcriptomic data. (E) Spatially resolved Grb14 expression in PD14, 1-month and 2-month mouse ovary respectively. A gradient of blue to red indicates the low to high gene expression level. (F) The expression patterns of Inhbb along trajectory (top) and dynamic expression patterns of follicle development associated genes along trajectory (bottom). SF, small follicle.

**Supplementary Figure S6.**
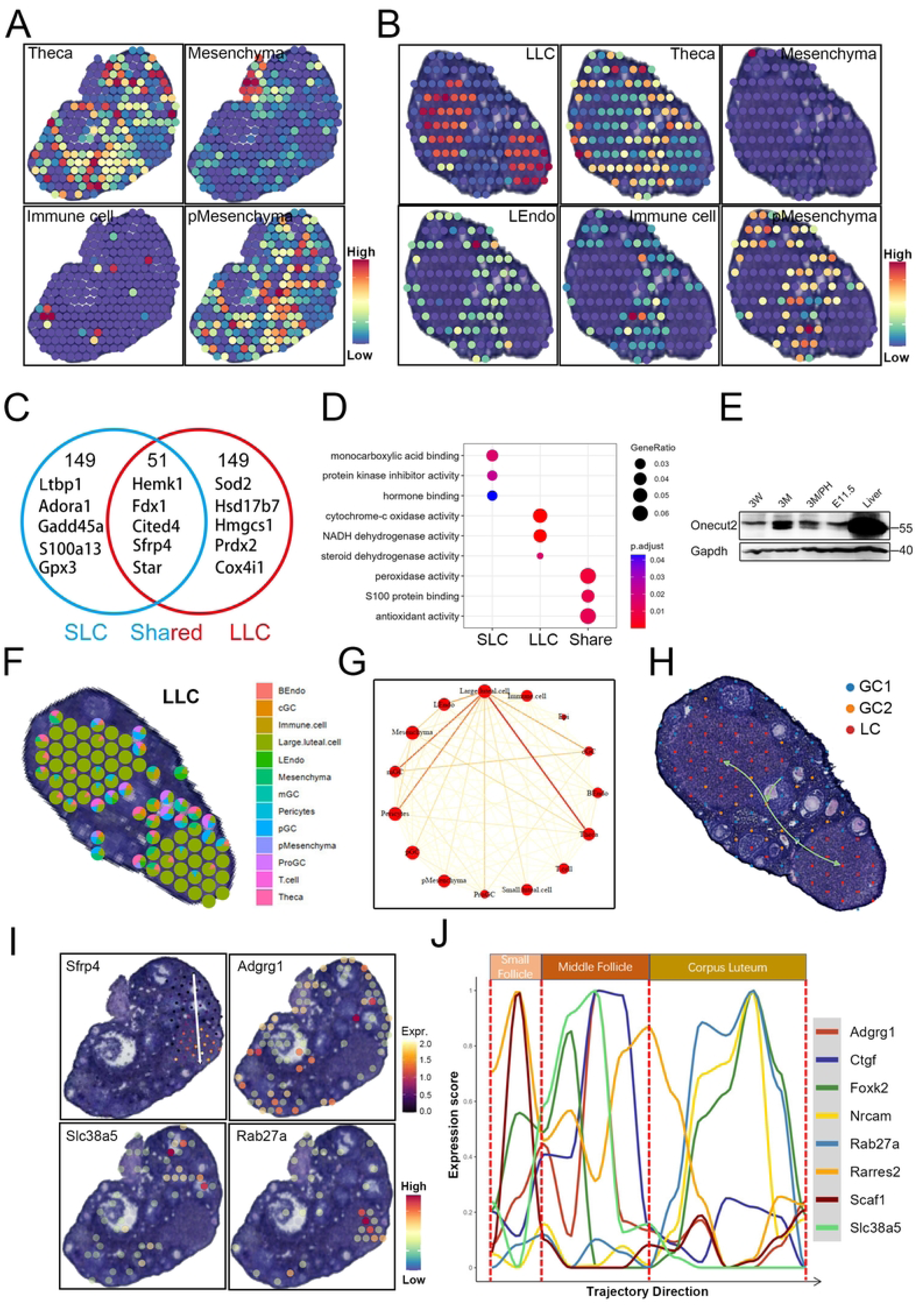
Identification of luteal cells and their specific spatial distributions. (A) Visualization of Theca cells, immune cells, mesenchyma cells and proliferate mesenchyma cells (pMesenchyma) on 2-month mouse ovary respectively based on integration of scRNA-seq cell type annotations with spatial transcriptomic data. (B) Visualization of large luteal cells (LLC), lymph associated endothelial cells (LEndo), Theca cells, immune cells, mesenchyma cells and proliferate mesenchyma cells (pMesenchyma) on another 2-month mouse ovary respectively based on integration of scRNA-seq cell type annotations with spatial transcriptomic data. (C) Venn diagram of top 200 differentially expressed genes in small luteal cells (SLC) and large luteal cells (LLC) demonstrates overlap between the gene populations in two subclusters of luteal cells. (D) The top GO terms enriched in the differentially expressed genes that unique to SLC (149), LLC (149) and shared genes (51) between LLC and SLC in Figure S6C. (E) The expression pattern of Onecut2 during ovary development. 3W, 3weeks; 3M, 3 months; 3M/PH, the ovary collected from 3-month mouse injected by pregnant mare serum and human chorionic gonadotropin; E11.5, embryonic day 11.5 ovary. (F) Spatial scatter pie plot representing the proportions of the LLC from the PD90 ovary reference atlas in the 2-month mouse ovary. (G) The spatial interactions among different cell clusters analyzed by SPOTlight. (H) Spatial trajectory showing the differentiation of granulosa cells in 2-month ovary section analyzed by stLearn. (I) The expression patterns of differentially expressed genes along trajectory. (J) Dynamic expression patterns of spatial trajectory (Fig. 6h) associated ordering genes along SPATA trajectory.

